# Accelerating drug development using zebrafish disease models supported by pharmacokinetic-pharmacodynamic modeling as new approach methodology (NAM)

**DOI:** 10.64898/2026.05.29.728504

**Authors:** Gabriel Forn-Cuní, Bart van Lieshout, Bjørn Koch, Cristina Villellas, Stijn Van Asten, Dina Berlina, Ellen Lanckacker, Bart Stoops, Rob J. Vreeken, Dirk Roymans, Elke H.J. Krekels, J.G. Coen van Hasselt, Herman P. Spaink, Rob C. van Wijk

## Abstract

The development of novel therapeutics for infectious diseases remains a global health priority. To accelerate the treatment development, innovative strategies through new approach methodology (NAM) are needed to bridge speed of *in vitro* with predictive power of *in vivo* studies, while reducing mammalian experiments. The zebrafish (*Danio rerio*), particularly the embryo/larva, has been established as a valuable non-mammalian *in vivo* model in biomedical research. We developed a standardized and streamlined workflow for the zebrafish as NAM, which consisted of 3 steps: drug selection and efficacy evaluation, internal exposure assessment, and PKPD modelling. Compounds with higher tolerated doses than minimum inhibitory concentration were selected. Drug efficacy was quantified through longitudinal individual fluorescence microscopy at baseline and 24 and 48h on treatment. Drug exposure was quantified in larval homogenates and exposure medium from 0-48h on treatment. The PKPD relationship was quantified by non-linear mixed effects modelling. For case study bedaquiline, PKPD was quantified using a one-compartment model with age-depending elimination, and an Emax concentration-response relationship on the delayed logistic bacterial growth function, with an EC50 of 26.5 µg/mL and an Emax of 1.09-1.37. In the case of clarithromycin, in contrast, negligible internal exposure after waterborne treatment was observed, illustrating the risk of false negatives without internal exposure assessments. Bactericidal efficacy was confirmed by intravenous drug injections, showing a clear dose dependent antimycobacterial effect. The standardized zebrafish NAM workflow presented here facilitates the translation of drug efficacy to higher vertebrates, reducing rodent studies to confirmatory or replacing them completely, thus accelerating drug development.

**Study Highlights:** *What is the current knowledge on the topic?:* There is an urgent need to accelerate development of anti-mycobacterial drugs while reducing preclinical rodent studies. The New Approach Methodologies (NAM) include non-mammalian *in vivo* experimental models, such as the phylogenetically lower vertebrate zebrafish (*Danio rerio*). Experiments in the zebrafish embryo (pre-hatching) or larva combine higher experimental throughput with lower ethical and resource constraints, while remaining predictive of higher vertebrates.

*What question did this study address?:* How can the zebrafish embryo/larva experimental model, supported by pharmacokinetic-pharmacodynamic (PKPD) modelling, be leveraged as New Approach Methodology (NAM) to accelerate drug development in infectious diseases by bridging *in vitro* throughput with *in vivo* predictability.

*What does this study add to our knowledge?:* We developed a standardized and streamlined workflow for the zebrafish as NAM, which consisted of 3 steps: drug selection and efficacy evaluation, internal exposure assessment, and PKPD modelling. Two anti-mycobacterial backbone drugs, bedaquiline and clarithromycin, were studied using the workflow. Bedaquiline dose-ranging PK and PD experiments supported quantification of the exposure-response relationship which approximated that in rodents. Clarithromycin, however, did not pass the workflow criteria, showing negligible internal exposure after waterborne treatment, illustrating the risk of false negatives without internal exposure assessments.

*How might this change clinical pharmacology or translational science?:* The use of zebrafish experiments supported by PKPD modelling as NAM shows significant promise in drug development in infectious diseases. Our developed experimental and computational workflow enables characterization of a drug’s *in vivo* exposure-response within approximately 2 months. With the exposure-response, further development can be supported by prioritizing or de-risking drug(s) and by informing dose and design of confirmatory mammalian experiments, to accelerate the development of the most promising drugs to patients.

## Introduction

The development of novel therapeutics for infectious diseases such as tuberculosis (TB) and non-tuberculous mycobacterial (NTM) infections caused by for example *Mycobacterium avium*, remains an urgent global health priority^1^. The traditional paradigm of drug development is lengthy, costly, and results in high attrition rates at the clinical development stage^2^. This is especially critical for the development of anti-mycobacterial therapies, where only a handful of drugs, such as bedaquiline and clarithromycin, has reached market approval in the last half century, and resistance against those is rising^3–6^. To accelerate the discovery and clinical translation of effective treatments, there is a clear need for innovative strategies that bridge the predictive power of *in vitro* and *in vivo* studies to patients, allowing for more efficient prioritization of drug candidates before late preclinical development and clinical trials^7^.

New approach methodologies (NAMs) are increasingly positioned as a pillar of modern drug development by complementing advanced experimental techniques with *in silico* approaches to improve translational success. In announcing its recent draft guidance on NAMs, the US Food and Drug Administration explicitly recognizes innovative methods ranging from complex *in vitro* and computational models to studies in non-mammalian, lower vertebrate organisms such as zebrafish (*Danio rerio*) as suitable contributors to nonclinical pharmacology packages when appropriately validated for context of use^8,9^. Such experimental systems gain maximal impact when coupled to quantitative, mechanistic modeling frameworks in model-informed drug development (MIDD). Pharmacokinetic-pharmacodynamic (PKPD) modelling has successfully been applied to predict clinical trial results for more than 10 anti-TB drugs from *in vivo* mouse experiments and translating those to clinical results^10–12^. Next, integrating innovative non-mammalian models with translational PKPD modelling will reduce reliance on traditional mammalian species while increasing the predictive value of preclinical decision making for clinical studies^13,14^.

The zebrafish^15^ has been established as a valuable animal model in biomedical research, toxicology screens, and drug development, with numerous examples of discovered or improved drugs supported by experiments in the zebrafish model^16^. Of special interest is the use of zebrafish during their embryo (pre-hatching) and larval developmental phase, offering a fast, tractable and informative whole individual *in vivo* model for studying disease mechanisms and testing therapeutic interventions. Given zebrafish’ high fecundity and the small size of their embryos and larvae, experiments can be performed at larger, faster experimental scale than with other vertebrates^13^. Especially when including automation using robotics, zebrafish disease models offer high-throughput experimentation with *in vivo* predictability. This is especially beneficial in anti-mycobacterial drug development, where testing combination regimens for treatment increases exponentially given all compounds currently under development^17^. In addition, zebrafish models are especially suited for the study and discovery of new drugs targeting infections affecting immunocompromised patients such as those caused by mycobacteria, because their developmental window allows for the study of the innate immune system without adaptive immunity^18^.

Quantification of drug efficacy (PD), as well as safety, is already commonly performed in zebrafish research, often exploiting their visual transparency for imaging-based measurements. When studying anti-infective efficacy, fluorescently labeled pathogens are utilized, which enables longitudinal experiments and offers a significant advantage in pharmacological investigations of infections, for example via high resolution repeated measurements within the same individual larva, and with high replicate numbers. This imaging process is easily automated using robotics. Quantification of drug’s internal exposure (PK), however, is often disregarded in zebrafish experiments. Bioactive molecules are dissolved in the incubation medium for waterborne treatment, and the internal concentration inside the larva is usually assumed to be the same as the external concentration. The dynamic PK processes of absorption, distribution, metabolism, and excretion are, thereby, ignored. To address the challenge of quantifying the PK in zebrafish larvae, we have previously developed an analytical workflow to quantify the internal drug exposure in homogenized zebrafish larvae for the characterization of the total internal amount as metric of internal exposure^19–21^. Despite the small size of the larvae, handling homogenates is straightforward, and it is possible to quantify internal exposure of a drug in individual larvae, as well as in pooled samples for less sensitive analytical methods. When drugs are administered through waterborne treatment, the larvae need to be washed to prevent contamination by skin adhesion. Sampling of internal exposure should be performed over time given that the organism is in development, with absorption and elimination processes maturing in a matter of days^19^. With quantitative exposure and response measures, a PKPD model can be developed to determine the efficacy (*E*_*max*_) and potency (*EC*_*50*_) of the drug of interest in zebrafish larvae. Leveraging translational factors for inter-species differences in pharmacological processes, or differences between pathogen strains in drug sensitivities, the PKPD relationship can be used to predict the drug concentrations required in higher vertebrates to achieve the desired drug effect.

To further develop the use of zebrafish as NAM in the context of translational drug development, our objective here is to formulate a standardized and streamlined workflow to study the PKPD of anti-infectives in zebrafish embryo/larva infection disease models, using standard-of-care drugs bedaquiline and clarithromycin treatment against the mycobacterial infections *M. marinum* and *M. avium* as case studies.

## Methods

### Workflow development

A standardized and streamlined workflow to study the PKPD of anti-infectives in zebrafish embryo/larva infection disease models upon waterborne treatment was developed, consisting of drug selection and efficacy determination through dose ranging PD, followed by internal exposure determination through dose ranging PK, concluded by exposure-response (PKPD) modelling. Drug selection is informed by solubility given waterborne treatment, toxicity, and drug sensitivity of the pathogen. Initial drug toxicity is assessed to calculate the maximum tolerated dose (MTD), and drug sensitivity is assessed by the minimum inhibitory concentration (MIC) *in vitro*. The first workflow criterium is formulated to only progress soluble drugs with an MIC lower than the MTD. From experience, the MTD can be reduced in infected larvae, which should be taken into consideration when comparing between the MIC and the MTD. Dose-ranging experiments are subsequently performed in zebrafish infected during the embryonic stage and followed-up during the larval stage, starting from the MTD and decreasing, to identify efficacious concentrations during the treatment window of 3 to 5 days post fertilization (dpf). Longitudinal fluorescence-based assays are used in blood island-infected embryos and larvae to assess drug efficacy. Manual infections are somewhat labor intensive, but automation through robotics has been successfully developed for both infections, and imaging^22,23^. The second workflow criterium is formulated to only progress drugs with clear efficacy. Following demonstration of *in vivo* drug efficacy, PK experiments, including washing in case of waterborne treatment, are designed and optimized. The third workflow criterium is formulated to achieve successful internal exposure determination, which can be challenging in case drug adhesion to the skin is high. Three experiments (∼1 month) follow to characterize internal exposure using mass spectrometry. Optionally, blood sampling can be performed to obtain volume of distribution as previously demonstrated^20,24^. Finally, mathematical models are applied to integrate PK and PD data, enabling the prediction of drug behavior to obtain *E*_*max*_ and *EC*_*50*_ *in vivo*, required to facilitate their translation to mammalian models and, ultimately, human clinical trials.

### Zebrafish husbandry

ABTL (wild-type) zebrafish were handled in compliance with the local animal welfare directives of Leiden University (License number 10,612) following standard zebrafish rearing protocols (https://zfin.org), which adhere to the international guidelines from the EU Animal Protection Direction 2010/63/EU. All experiments were performed on zebrafish embryos or larvae up to 5 dpf, which have not yet reached the free-feeding stage. Embryos were collected in egg water (Sera Marin salt, 05420; 60 µg/ml in distilled deionized water) and maintained in E3-buffered medium (300 mM NaCl, 10.2 mM KCl, 19.8 mM CaCl_2_-2 H_2_O, 19.8 mM MgSO_4_-7H_2_O in egg water) at 28.5°C for the duration of the experiments. Prior to infections and imaging, embryos and larvae were anaesthetized with 0.02% buffered 3-aminobenzoic acid ethyl ester (tricaine; Sigma-Aldrich, A-5040) and positioned on a 2% agarose plate in E3.

### Selection of drugs and efficacy determination

For the development of the workflow, the non-tuberculous mycobacterial infections caused by *M. marinum* and *M. avium complex* were selected as case studies. The backbone drugs of current successful regimens against mycobacterial infections, bedaquiline and clarithromycin, were selected for both infection case studies. The zebrafish infection model by *M. marinum* has been established as a predictive model for tuberculosis in higher vertebrates, and is therefore assessed in most detail for predictive purposes. The MTD was determined using the Fish Embryo Acute Toxicity Test (FET; OECD Test Guideline 236), and the MIC was determined according to EUCAST reference protocol for MIC determination using broth microdilution in Middlebrook 7H9 (**Supplementary methods**).

Infection experiments with *Mycobacterium marinum* (Mm) or *Mycobacterium avium complex 101* (MAC101) expressing the fluorescence protein mWasabi were performed according to standard procedures^24,25^, **Supplementary methods**). After infection, larvae were individually kept in 48-well plates with 500 μL of medium containing the tested drug for waterborne treatment. Only if waterborne treatment was unsuccessful, was direct injection performed at 3 dpf in the bloodstream via the duct of Cuvier. For assessment of bacterial burden, larvae were positioned on a 2% agarose plate and imaged using a Leica M205 FA fluorescence stereomicroscope equipped with a DFC345 FX monochrome camera. Bacterial burden was determined longitudinally based on fluorescent pixel quantification using QuantiFish ^26^.

### Internal exposure determination

For drug adhesion and washing efficiency experiments, larvae were submerged at the highest drug concentration for 1 minute and washed 5 times with 2 mL of 20/80 methanol/water solution (v/v). For determination of internal drug exposure and absorption and elimination dynamics, larvae were submerged at 0.25, 0.5, 1, 5, and 10X MIC concentrations to consider possible non-linear kinetics including saturation. For the main concentration (1X MIC), sampling was performed at 0, 0.5, 1, 2, 4, 8, 24, 30, and 48 h post incubation after the start of treatment (3 dpf). For the rest of the concentrations, samples were taken more sparsely at 4, 24, 30, and 48 h post incubation after the start of treatment, for later integrative modelling across concentrations and timepoints. For the determination of the elimination rate and absolute clearance, larvae were treated at the 0.25, 1X, and 10X MIC drug dosage for 48 h and then transferred to non-drug containing E3-buffered medium. Samples were taken after 0, 0.5, 1, 2, 4, and 6 h after stopping the waterborne treatment. All experiments were performed in triplicate. Drug dissolved in E3-buffered medium stability was also assessed (**Supplementary methods**). Internal drug exposure in zebrafish larvae was quantified as drug amounts in larval homogenates similarly as previously described^19–21,24^, **Supplementary methods**)

### Exposure-response modelling

Upon successful waterborne treatment, (non)linear mixed effects PKPD modelling was performed by a sequential approach in which the PK model was built first and then appended with the PD component to develop the final PKPD model, using fixed PK estimates to describe the exposure of the infection experiment. In case of failing workflow criterium three, i.e. failure to achieve successful internal exposure determination, no PKPD modelling was performed, for example because waterborne treatment did not yield sufficient internal exposure for observable PK and subsequently PD observations.

PK modelling was performed simultaneously on the drug measurements in the incubation medium (ng/mL) and zebrafish larvae (ng/larvae). Here, a one-compartment PK model with first-order absorption and elimination was assumed for the zebrafish because only whole-body homogenates were measured^20^. Additionally, residual unexplained variability (RUV) was the only measure of variability used to describe the PK data, as inter-individual variability (IIV) could not be quantified using non-longitudinal data from destructive sampling. The drug concentration in the medium was modelled as a separate compartment with a first-order degradation rate to account for drug degradation. Separate additive, proportional, or combined additive and proportional error models were tested for medium or zebrafish observations. To account for possible developmental changes in the PK in zebrafish larvae^21^, stepwise covariate modelling of age (dpf) on the absorption rate constant or the elimination rate constant was performed using additive (Eq. 1), linear (Eq. 2), exponential (Eq. 3), or power relationships (Eq. 4). Herein, the covariate effect on the population parameter (*θ*_*pop*_) was defined through the relationship (*θ*_*cov*_) of an individual’s covariate value (*COV*_*i*_) and the covariate’s reference value (*COV*_*ref*_= {4 *dpf*, 5 *dpf*}). These were used to calculate the individual parameter value (*P*_*i*_). For better translatability, internal exposure measures were expressed as concentrations, rather than total amounts, using volumetric measurements of the larvae as the volume of distribution^27^. For the medium, stepwise dose correcting factors (DCF) were assessed to account for differences between the expected and measured initial drug concentration in the medium.

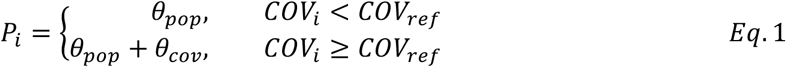

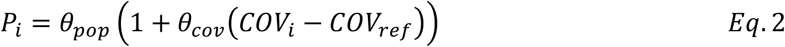

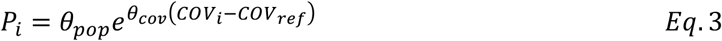

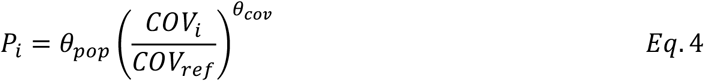

The PD was modelled on the bacterial burden by using log_10_ transformed measurements of the emitted fluorescence, which was calibrated with colony-forming units (**Figure S1**). Different structural models were applied to the data to describe the changes in the bacterial burden 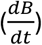 as described before^24^. Firstly, exponential (Eq. 5), Gompertz (Eq. 6), and logistic (Eq. 7) equations were considered, which describe the bacterial growth rate (*GROWTH*) with a growth rate constant (*k*_g_) that is modulated by the ratio between the bacterial burden (*B*) and the maximum bacterial capacity (*K*).

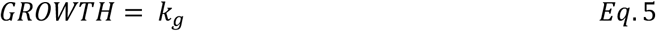

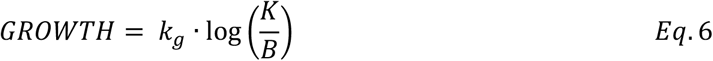

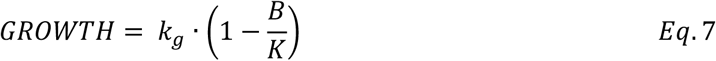

The relationship between the drug effect size (*EFF*) and the drug concentration (*C*_*bqq*_) was modelled with linear (Eq. 8), *E*_*max*_, (Eq. 9) or sigmoid *E*_*max*_ (Eq. 10) relationships. For the linear relationship the effect increases proportionally to the concentration with *SLP* and for the *E*_*max*_ equations increase in effect was saturable to a maximum of *EXAX*, with half the maximum effect at a concentration of *EC*50, and a steepness factor of *γ*.

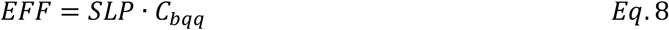

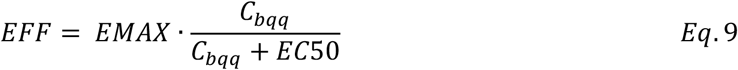

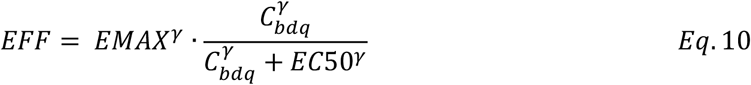

The effect itself was parameterized as either proportional to the growth rate constant (Eq. 11) or as a first-order killing rate constant on the bacterial population (Eq. 12).

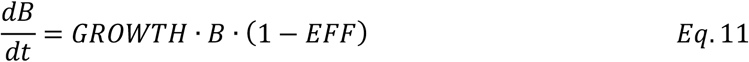

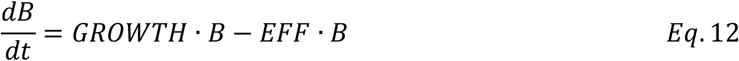

IIV with a log-normal distribution was evaluated on PD parameters, informed by the repeated longitudinal PD observations per individual larva, and *a priori* on the inoculum size. Lastly, age-dependent changes in PD were included in the model through either the inclusion of a stepwise decrease in the growth rate constant, a stepwise increase in the maximum effect, an effect compartment, or a timed delay in growth after inoculation (Eqs. 1-4). Variability was included through an additive error model, equivalent to an exponential error model on untransformed data.

Model selection was based on numerical, graphical, and pharmacological criteria. Numerically, the likelihood ratio test of the difference in objective function value, assuming a Chi-squared distribution, was utilized, as well as parameter precision through relative standard errors of the estimates (RSE) below 50%. Graphically, goodness-of-fit plots and visual predictive checks (with *n* = 500 simulations) were utilized. Finally, the physiological and mechanistic plausibility of parameter estimates was taken into consideration. Modelling was performed using *NONMEM* v7.5.1^28^, supported by *Pirana* v23.10.1^29^ and *PsN* v5.3.1^30^. The first-order conditional estimation method with interaction was used in NONMEM^28^. For dose-response statistics, log10-transformed bacterial burden data were compared between treatment groups using the Kruskal– Wallis test, followed by Dunn’s post-hoc pairwise comparisons of each drug dose against the vehicle control (DMSO for waterborne treatment, PBS for injection). Data processing and graphical analysis was performed with *R* v4.3.3^31^ using the integrated development environment *Rstudio* v2023.12.1+402 (RStudio, PBC., Boston, MA). Visual model assessments were generated with the *vpc* v1.2.2 package^32^.

## Results

A streamlined workflow was developed for conducting PK and PD studies in zebrafish embryos and larvae, enabling the rapid pharmacological evaluation of novel antimycobacterial drugs (**Figure 1**). This workflow involves sequential steps, experimentally starting with determining drug efficacy through comparison of MIC to MTD and (automated) PD evaluation, followed by PK studies which are more cumbersome in zebrafish larvae and therefore only applied to the most promising compounds, and ending with PKPD modelling of the exposure-response relationship. Multiple workflow criteria were formulated, allowing for efficient screening and prioritization of promising drug compounds. By leveraging this pipeline, *in vivo EC*_*50*_ and *E*_*max*_ values can be generated within a short timeline of approximately 2 months, thereby reducing the number of (mammalian) animals required for subsequent confirmatory studies.

**Figure 1.**
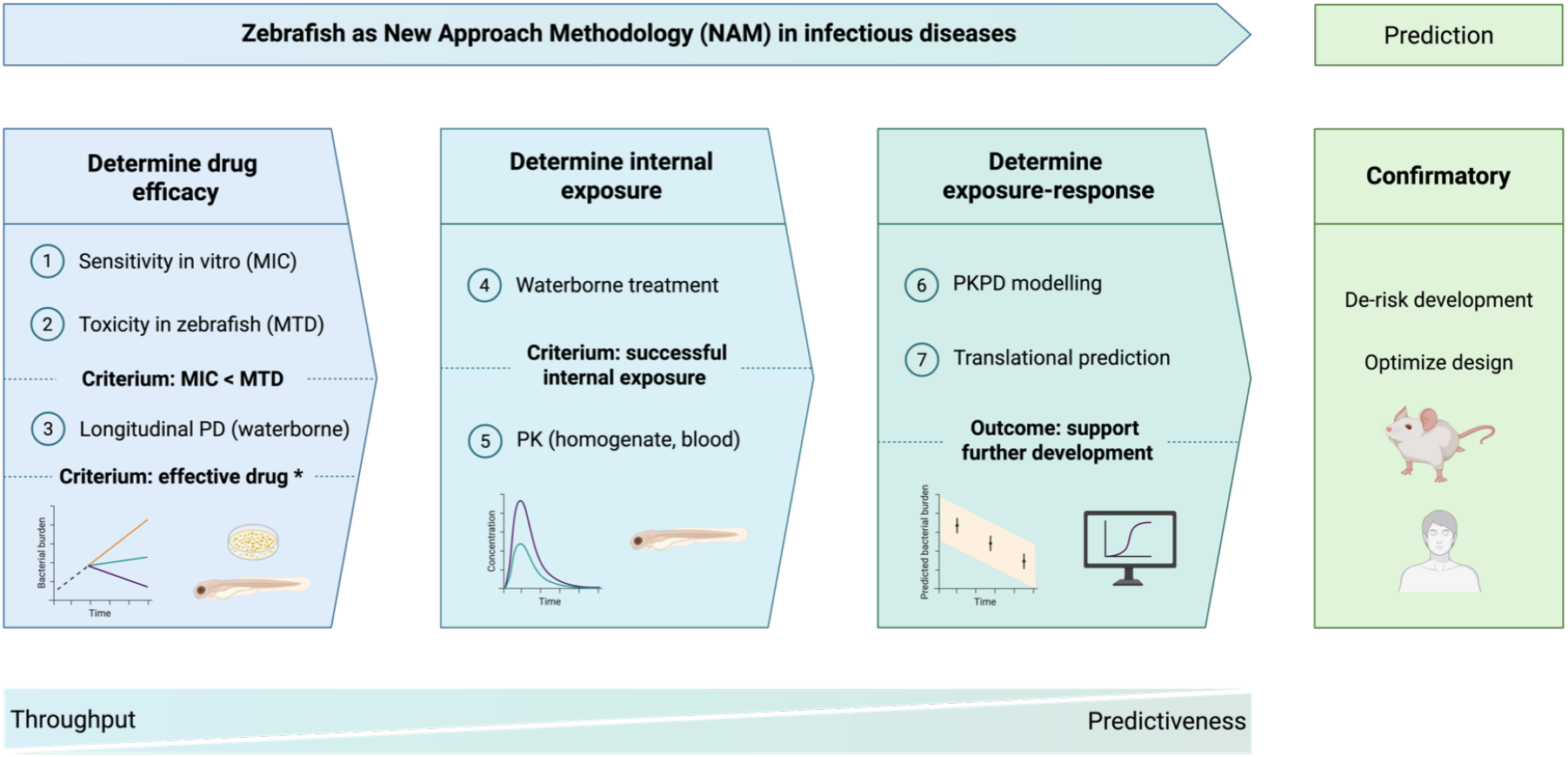
Standardized and streamlined workflow to study anti-infectives pharmacokinetics (PK) and pharmacodynamics (PD) in zebrafish embryo/larva infection disease models. The first pillar focusses on determining drug efficacy (PD), informed by the maximum tolerated dose (MTD) related to the minimum inhibitory concentration (MIC) of the drug (steps 1–3). The second pillar has a focus on determining internal exposure (PK), quantified in zebrafish larvae using sensitive liquid chromatography coupled to mass spectrometry (LC/MS) (steps 4–5). The third pillar has a focus on determining the exposure response through PKPD modelling, supporting further development with confirmatory mammalian or clinical studies by translational predictions from zebrafish larvae to higher vertebrates (steps 6–7). Each pillar closes with a criterium that has to be met before progressing to the next. *A compound without an effect after waterborne treatment is not necessarily inefficacious, as insufficient internal exposure may prevent an effect from being observed. Such compounds can be administered through an alternative route, such as intravenous injection, and re-enter the workflow at step 3. Image created in BioRender. Forn-Cuni, G. (2026) https://BioRender.com/nn7nbzb

The streamlined workflow was performed as case studies for bedaquiline and clarithromycin in the treatment of mycobacterial infections. Both compounds showed different characteristics and outcomes in the workflow, which will be discussed in detail below.

### Bedaquiline: drug efficacy

For the case study of bedaquiline in treatment of *M. marinum*, the MIC and MTD were compared first. The MIC of bedaquiline to *M. marinum* is 0.1 mg/L (0.18 µM)^33^, and has generally been shown to be lower than 0.25 mg/L (0.45 µM) for several mycobacteria, including *M. tuberculosis*^34–36^, as well as for slow-growing and fast-growing mycobacteria isolates^37–39^. Within the MTD assay using waterborne bedaquiline treatment of zebrafish, no toxicity was found up to the maximum concentration tested of 50 µM. With the favorable comparison (>25-fold) between MIC and MTD, the next step was to determine the dose-ranging PD. The PD analysis was performed on a total of 60 zebrafish larvae with longitudinal measurements of their bacterial burden in fluorescence, encompassing at least n=5 across the timepoints and doses of bedaquiline (0, 0.5, 1, 5, 10, and 50 µM). Comparing the difference in bacterial burden between measurements at 0 and 48 hours post-infection, a postponed growth by *M. marinum* after inoculation could be seen (**Figure 2A**). Furthermore, a clear dose-response relationship was observed when accounting for experimental variability in starting inocula. Between the third and fourth day of infection, there was a decrease in the rate of change in the bacterial burden, irrespective of the dose or bacterial burden. Separately, a significant dose-dependent effect on *M. avium* burden was observed (**Figure S2**).

**Figure 2.**
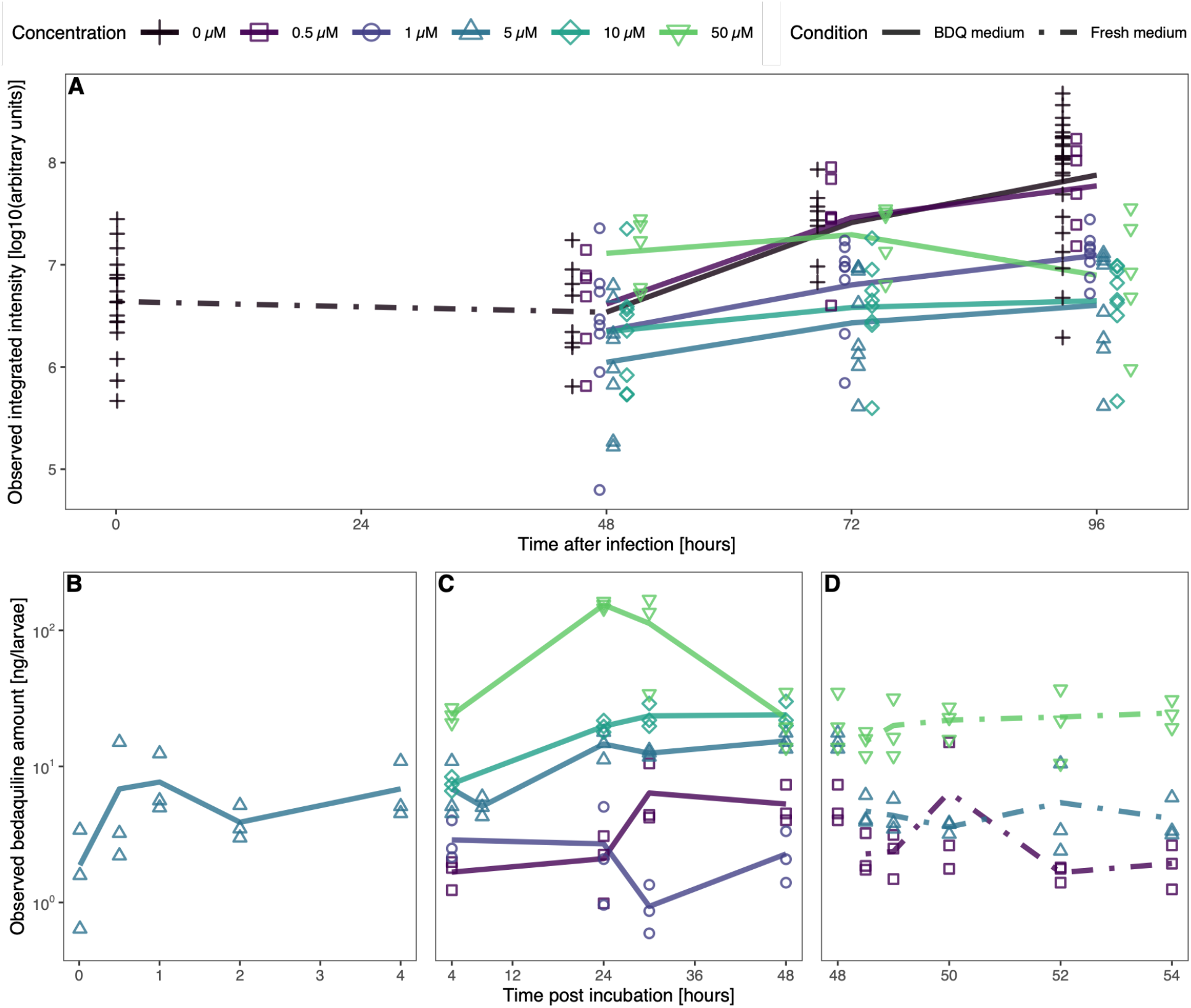
Measured drug efficacy (top) and quantities of bedaquiline per zebrafish larvae (bottom) across different levels of exposure to bedaquiline in the medium during incubation from 3 days post infection (dpf), with bedaquiline removed from after 48 hours of incubation. The infected larvae were exposed to a range of bedaquiline concentrations from 48 hours post infection. Observed changes over time in the bacterial burden of zebrafish larvae infected with *Mycobacterium marinum* shown as the log10 values of the integrated intensity (A). Lines connect median values of observations which are shown as symbols. Highlighted for the drug exposure are the absorption (B), steady state (C), and elimination (D) phases.

### Bedaquiline: internal exposure

The next step was to determine bedaquiline PK. A total of 45 measurements of bedaquiline concentrations in the medium were taken over time for doses of 0.5, 5, and 50 µM. Furthermore, the amount of bedaquiline in ng was quantified in 120 homogenates of n=5 larvae that were dosed with either 0.5, 1, 5, 10, or 50 µM. Both measurements were obtained for n=3 replicates at each timepoint. The PK profile of bedaquiline in the zebrafish larvae can be described to have a rapid absorption phase (**Figure 2B**) and a dose-dependent steady state (**Figure 2C**). After the removal of bedaquiline from the medium at 48 hours post incubation, an initial decrease in the internal exposure could be observed followed by a shallower decrease consistent with bedaquiline’s long half-life (**Figure 2D**).

### Bedaquiline: exposure-response modelling

PKPD modelling was performed on the bedaquiline data. The final PK parameters for bedaquiline in zebrafish larvae were 0.000941 h^-1^ for the absorption rate constant and 0.089 h^-1^ for the elimination rate constant, which increased by 0.122 h^-1^ at 5 dpf as shown in the covariate analysis (*p*=0.025, **Table 1**). Differences between the expected and measured initial drug concentration of the medium were addressed through the addition of an initial DCF for doses above 5 µM (*p*<<0.001), followed by a second DCF for doses above 10 µM (*p*<0.001). All parameters of the PK model were estimated with RSEs below the 50% threshold.

**Table 1.** Parameter estimates for the pharmacokinetic and pharmacodynamic components of the model.

| Parameter (unit) | Description | Estimate (RSE%)<br>[% shrinkage] |
| --- | --- | --- |
| *Structural and covariate parameters* |  |  |
| $k_{10}$ ($\text{h}^{-1}$) | Degradation rate constant of bedaquiline in medium | 0.0189 (30.6) |
| $k_{12}$ ($\text{h}^{-1}$) | Absorption rate constant of bedaquiline by zebrafish larvae | 0.000941 (40.6) |
| $k_{21}$ ($\text{h}^{-1}$) | Elimination rate constant of bedaquiline by zebrafish larvae | 0.089 (33.7) |
| $k_{21,5dpf}$ ($\text{h}^{-1}$) | Increase in elimination rate constant at 5 dpf | 0.122 (44.6) |
| $DCF_{0.5 < x \leq 10}$ (-) | Ratio dose correcting factor for doses ranging from 1 to 10 $\mu\text{M}$ | 0.299 (28.4) |
| $DCF_{10 < x}$ (-) | Ratio dose correcting factor for doses above 10 $\mu\text{M}$ | 0.216 (26.9) |
| $B_0$ ($\log_{10}(\text{a.u.})$) | Bacterial inoculum size | 6.52 (1.7) |
| $k_g$ ($\text{h}^{-1}$) | Bacterial growth rate constant | 0.0853 (26) |
| $t_{delay}$ ($\text{h}^{-1}$) | Time delay in bacterial growth | 48 (20.4) |
| $K$ ($\log_{10}(\text{a.u.})$) | Zebrafish carrying capacity | 8.19 (0.9) |
| $EMAX$ ($\text{h}^{-1}$) | Ratio reduction in growth rate by bedaquiline | 1.09 (42.9) |
| $EMAX_{3dpi}$ (-) | Ratio reduction in growth rate by bedaquiline after 3 dpi | 1.37 (14.5) |
| $EC50$ ($\mu\text{g/mL}$) | Bedaquiline concentration for 50% of the maximum response | 26.5 (47.2) |
| *Statistical parameters* |  |  |
| $RUV_{prop}^{med}$ ($\sigma^2$) | Proportional residual unexplained variability of bedaquiline in medium | 0.381 (38.3) [1] |
| $RUV_{add}^{zf}$ ($\sigma^2$) | Additive residual unexplained variability of ng bedaquiline in zebrafish | 11.1 (10.4) [0.1] |
| $RUV_{prop}^{zf}$ ($\sigma^2$) | Proportional residual unexplained variability of bedaquiline in zebrafish | 0.247 (36.8) [0.1] |
| $RUV_{add}^{rfu}$ ($\sigma^2$) | Additive residual unexplained variability of $\log_{10}$ bacterial fluorescence | 0.0443 (19) [31] |
| $IIV_{inoc}^{rfu}$ (CV%)<sup>a</sup> | Inter-individual variability of the inoculum sizes | 186.6 (14) [5] |
| $IIV_{growth}^{rfu}$ (CV%)<sup>a</sup> | Inter-individual variability of the growth rate constants | 74.6 (20) [27] |

Abbreviations: h, hour; dpf, days post fertilization; a.u., arbitrary units; CV, coefficient of variation.
<sup>a</sup> The coefficient of variation for the inter-individual variability (IIV) was calculated with $CV\% = 100\% \cdot \sqrt{e^{\omega^2} - 1}$ .

Bedaquiline PD model fitting was performed with the bacterial burden data that involved the omission of one replicate in the 0 µM dose group due to negligible growth. The final PD model consisted of a delayed logistic growth function, an *E*_*max*_ concentration-response relationship between bedaquiline concentration and effect, an effect parameterized as proportional to the growth rate constant of *M. marinum*, and inter-individual variability on the inoculum and growth rate constant. A sigmoid *E*_*max*_ relationship yielded a statistically significant improvement of the model fit, but with unacceptable imprecision in parameter estimates. The model improved statistically significantly with an increase in the *E*_*max*_ after 3 dpi (*p*<0.01). Primary estimated PD parameters were a growth rate constant of 0.0853 h^-1^, an *E*_*max*_ of 1.09 times the growth rate constant before 3 dpi and 1.37 afterwards, and an *EC*_*50*_ of 26.5 µg/mL, each with an RSE below 50% (**Table 1**).

The final model PKPD structure can be seen in **Figure 3**. Visual assessment of the final model fit through visual predictive checks highlights that the characteristics in the observations are captured well (**Figure 4, Figure S3**). However, certain dose groups showed shifted prediction intervals due to the omission of structural dose group-level variability in the inoculum size in the final model. These deviations ultimately did not affect the model’s description of the individual data (**Figure S4**). The high additive error for zebrafish larvae internal exposure, seen mainly at 0.5 and 1 µM doses, may have been due to a rapid absorption with uncertainty in the timing of early datapoints to inform accurate measures of residual error. The goodness-of-fit plots indicate that there were no major concerns with the residuals, aside from the aforementioned points (**Figure S5**).

**Figure 3.**
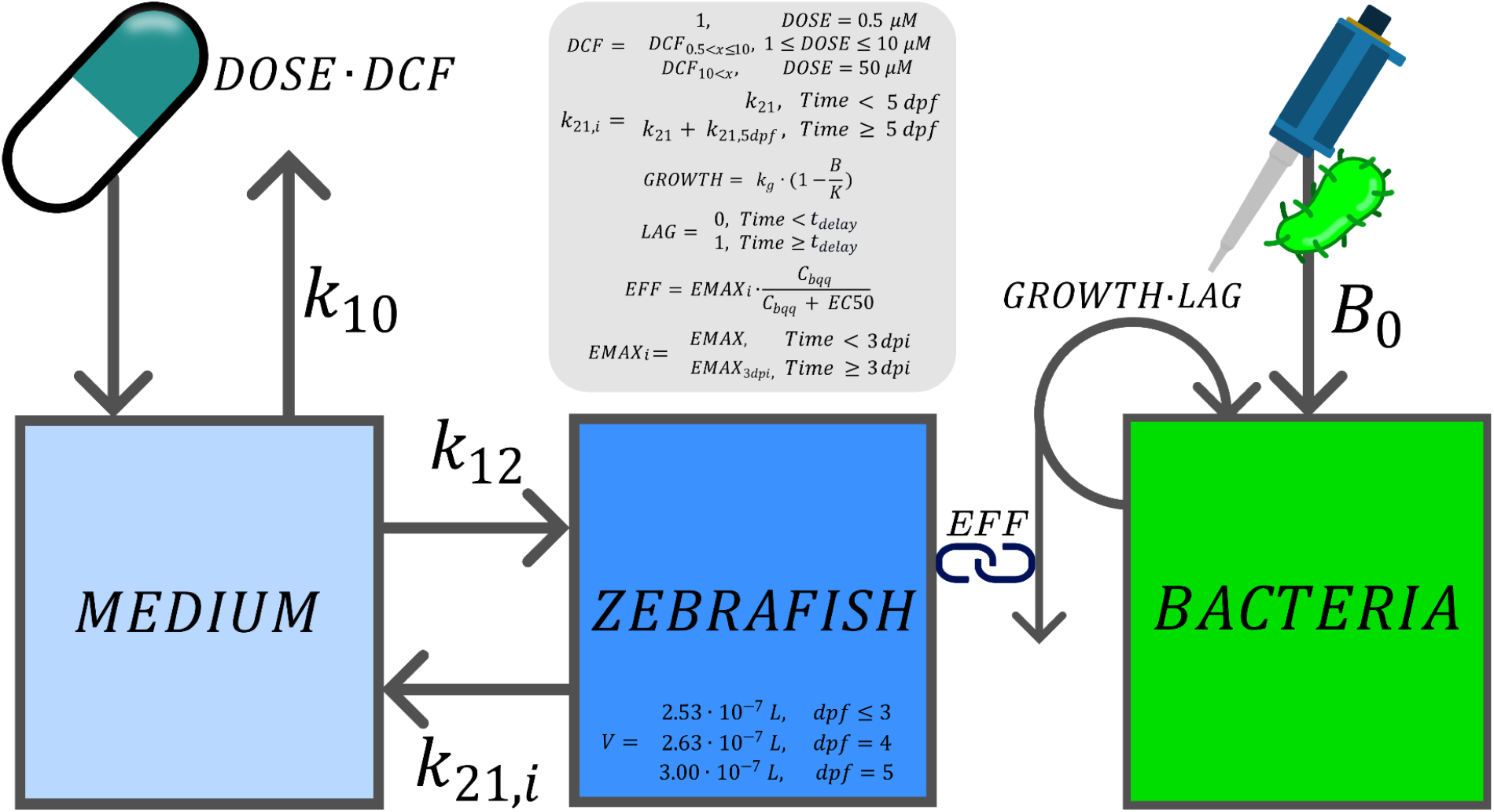
Final pharmacokinetic-pharmacodynamic model overview for bedaquiline in zebrafish larvae. Abbreviations and parameter descriptions are given in **Table 1**.

**Figure 4.**
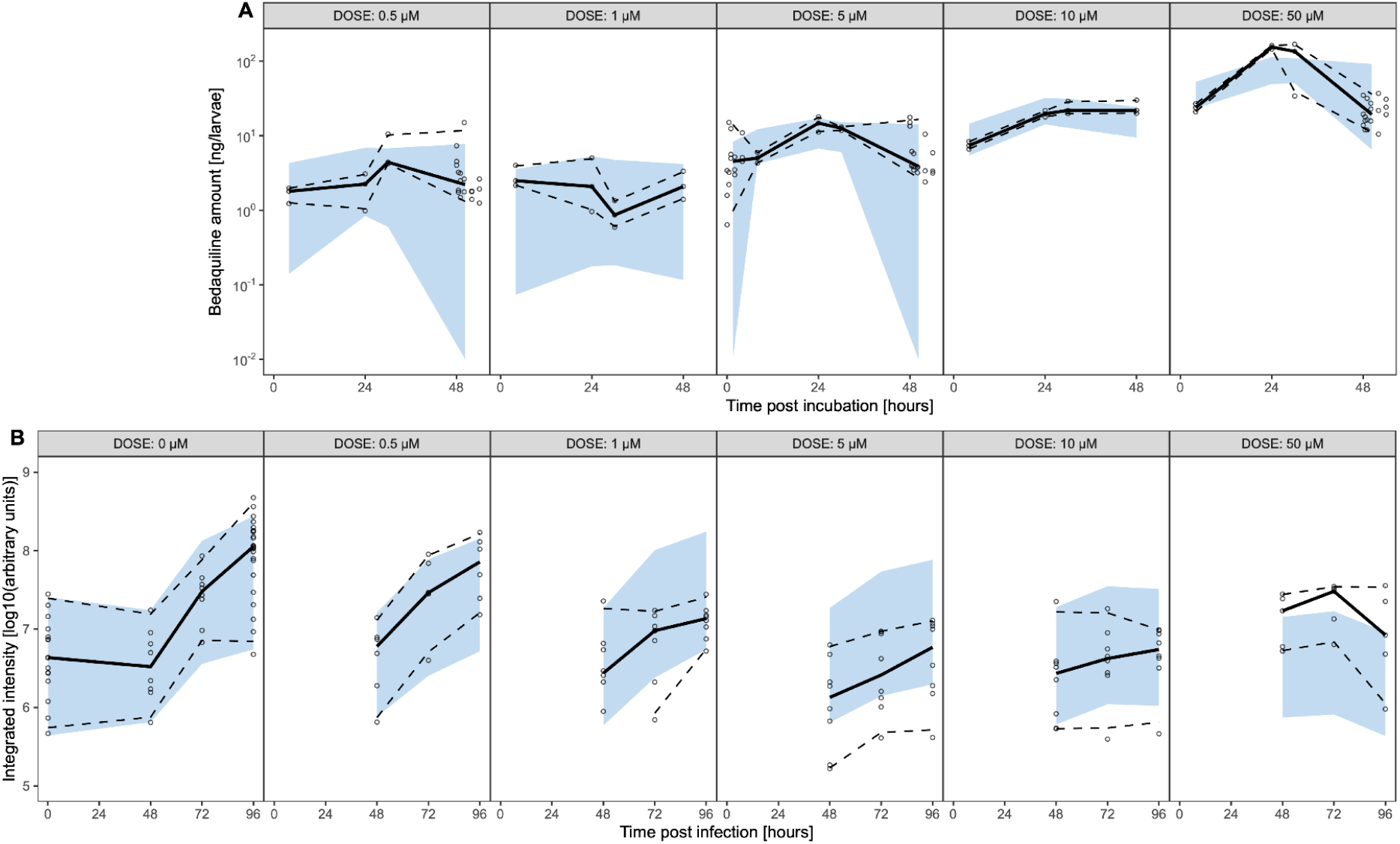
Visual predictive checks of simulations. (*n* = 500) that were performed with the pharmacokinetic (A) and pharmacodynamic (B) model fits to produce predictions of the total internal exposure to bedaquiline of zebrafish larvae and the bacterial burden in integrated intensity, respectively. Shown are the individual observations (open circles), the median (solid black line), and the 95% quantile (dashed black) along the 95% prediction interval (blue ribbon). Negative predicted values of bedaquiline in zebrafish larvae were censored to 0.01 ng/larvae.

### Clarithromycin did not pass workflow criteria

Similar to the bedaquiline case study, the MIC of clarithromycin was higher than its MTD in zebrafish. However, clarithromycin showed negligible internal exposure after waterborne treatment (**Figure S6**), consistent with the only previous report of waterborne clarithromycin efficacy in zebrafish larvae, in which an effect on *M. abscessus* infection required 170-fold the *in vitro* MIC of 4 µM, corresponding to 680 µM in the exposure medium, without internal exposure being determined^40^. Clarithromycin concentrations in exposure medium were stable, and washing was optimized to prevent skin adhesion from contaminating the internal exposure samples. Still, the internal exposures were less than 0.1 % of the external medium, and apparent dose dependency in exposure may have been caused more by insufficient washing than internal exposure differences. Therefore, clarithromycin did not pass the criterium of successful internal exposure. To confirm clarithromycin’s bactericidal efficacy, direct intravenous compound injections were performed. A clear dose dependent effect on both *M. marinum* and *M. avium* was shown (**Figure 5**). Clarithromycin thus represents a false-negative case in the proposed workflow, and PKPD modelling was not performed given criteria not being passed of this study which focused on streamlined workflow based on waterborne exposure. Internal exposure was not quantified under intravenous administration, so the injection experiment establishes efficacy across 2 dose levels, but not an exposure-response relationship for this compound.

**Figure 5.**
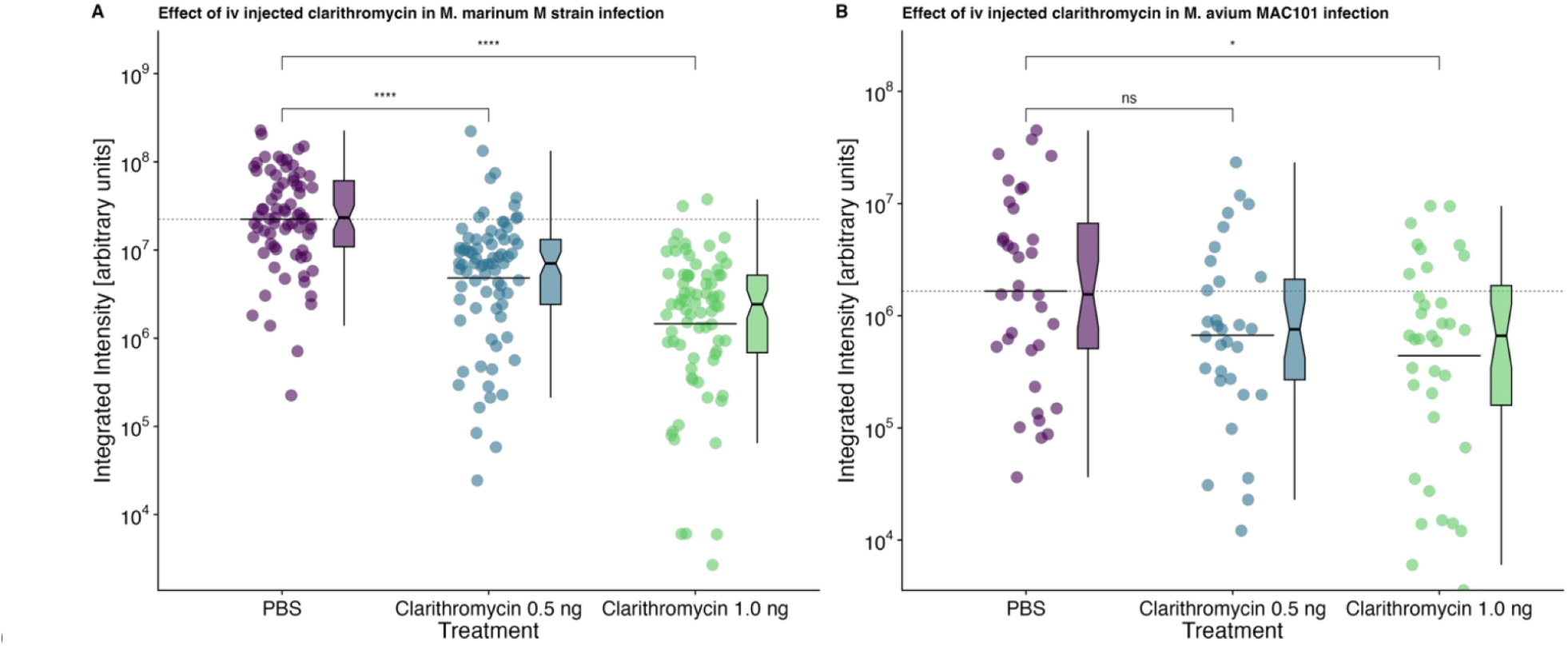
Clarithromycin is effective against *Mycobacterium marinum* and *Mycobacterium avium* MAC101 strains when intravenously injected. Integrated intensity of the fluorescence signal (symbols, boxplot) is shown for two dose groups of 0.5 and 1.0 ng injected clarithromycin, as well as phosphate-buffered saline (PBS) injection control group. Groups were compared by Kruskal–Wallis with Dunn’s post-hoc comparisons against the PBS control: Ns = not significant, * p < 0.05, ** p < 0.01, *** p < 0.005, **** p < 0.001.

## Discussion and conclusion

The zebrafish supported by PKPD modelling, can support drug discovery and development as NAM, prioritizing and de-risking drugs for further development in mammalian experiments and clinical trials. To that aim, a streamlined workflow is presented here, integrating experimental and computational techniques, applied to mycobacterial infections with two case studies. The workflow resulted in quantification of *in vivo* drug efficacy and potency that are within 1-2 orders of magnitude of mammalian values in ∼2 months^41^. The pharmacology of bedaquiline in the zebrafish infection disease model was successfully quantified, for both internal exposure over time as well as the exposure-response relationship. The workflow leverages fluorescence microscopy to quantify the PD, which is easy and fast to measure in comparison to culture-based measurements. PK is more challenging to quantify given the small size of the organism. Therefore, PKPD analysis should be prioritized for promising compounds as observed in the high-throughput assays, with the risk of missing false negative compounds such as the clarithromycin case study that do not reach efficacious internal exposures.

The quantification of PK and PD relies on several assumptions. Internal exposure is quantified in larval homogenates, resulting in drug amounts per larva in comparison to blood or plasma concentrations utilized in later preclinical drug development and clinical practice. Here, the assumption is made that drugs distribute homogenously throughout the organism, equating the volume of distribution to the total body volume as determined earlier by three-dimensional tomographic microscopy^27^. A blood sampling technique from 5 dpf zebrafish larvae has been developed previously^24^ but is labor intensive and requires a more sensitive analytical method for bedaquiline than available. However, for paracetamol (acetaminophen) and for isoniazid, the assumption on total body volume proved to be within 30% and 10% of the volume of distribution determined through blood sampling, respectively^20,24^. In addition, internal exposure quantified in homogenate represents the total amount in the organism and does not resolve its distribution between tissues or cellular compartments, so preferential accumulation at the site of infection cannot be excluded.

Bedaquiline clearance through metabolism through CYP3A4 as major pathway is suggested clinically to mature with age^42,43^. This maturation was captured in the zebrafish larva model between 4 and 5 dpf, as has been shown for other metabolic pathways in zebrafish^21,44^. The short duration of the PK studies in zebrafish larva does not allow for full metabolic maturation characterization, but as the focus was on quantifying internal exposure in zebrafish during the experimental period to quantify the PKPD relationship, characterizing full metabolic maturation is outside the scope of this work. Microbiologically, the PKPD of bedaquiline can vary for different bacterial strains, but it can be assumed to scale proportionally to their minimum inhibitory concentration (MIC) which are determined more readily and can be utilized as translational scaling factor^24,45^. The accuracy of PKPD projections is also dependent on other translational factors such as plasma protein binding, acute versus chronic infections, mycobacterial sub-populations, and differences in body temperature between zebrafish disease models and humans.

Despite its utility in preclinical screening, the zebrafish model presents inherent limitations that can impact drug selection and prioritization. While zebrafish larvae offer a cost-effective and high-throughput system for assessing early bactericidal activity, they lack several key mammalian physiological features, such as complex lung architecture and adaptive immunity. Waterborne treatment can result in false negatives, where drugs with potential efficacy in humans may be overlooked due to limited internal exposure during the experiments in zebrafish larva. This risk can be mitigated by changing the route of administration, as was done here for clarithromycin, where intravenous injection confirmed the bactericidal activity absent on waterborne treatment. Because injection has a considerably lower throughput than waterborne treatment, it is best applied selectively to promising compounds most likely to be false rather than true negatives (i.e., a very low MIC, physicochemical properties associated with poor uptake, such as high molecular weight and ionizability). A compound that fails on internal exposure is then classified as not evaluable by waterborne treatment rather than as inefficacious, and is escalated rather than discarded. Such limitations highlight the need for MIDD and computational models informed by complementary datasets.

Nevertheless, the zebrafish infection model remains a valuable tool for accelerating anti-mycobacterial drug development by enabling rapid screening of drug efficacy in a whole-organism context. Unlike *in vitro* assays, zebrafish models provide insight into the kinetics and dynamics of drug absorption and disposition, and the impact of host immune responses on bacterial clearance. The transparency of zebrafish larvae allows real-time visualization of bacterial burden and granuloma formation, facilitating dynamic assessments of drug action. As such, the zebrafish infection disease model combines experimental throughput at *in vitro* rates with whole organism predictiveness of *in vivo* studies^13,46^. Additionally, the experimental throughput enables testing drug combinations in a scalable manner in the future, enhancing regimen selection by identifying synergistic interactions early in the development pipeline. The fact that zebrafish larvae represent a complete organism also allows for the testing of host-directed therapies, providing a comprehensive understanding of disease mechanisms and treatment efficacy^25,47^. When integrated with PKPD modeling, results from zebrafish studies can optimize experimental design regarding the choice of drug and refine experimental design for mammalian models, thereby improving translational efficiency^14,48,49^.

The use of the zebrafish in anti-infective drug development shows significant promise, though further refinement is required. The established experimental methodologies and computational approaches shown here facilitate the translational scaling of drug efficacy findings, for example in guiding initial dose selection for mammalian models. The ability to generate in early drug development high-throughput *in vivo* pharmacological data that can be quantitatively translated to higher vertebrates, has the potential to shift rodent studies from exploratory to confirmatory experiments, offering substantial advantages in terms of efficiency, cost reduction, and ethical constraints, thus acceleration of drug development.

## Acknowledgements

This project has received funding from the Innovative Health Initiative 2 Joint Undertaking (JU) under grant agreement No 853903 (RESPIRI-TB) and No 853932 (RESPIRI-NTM). This Joint Undertaking receives support from the European Union’s Horizon 2020 research and innovation programme and EFPIA. We gratefully acknowledge the support of the Farmaceutische Wetenschappen Sectorplan, a component of the Sectorplan Bèta-II initiated by the Ministry of Education, Culture and Science of the Netherlands.

## Data availability

The final dataset and model code have been deposited at **10.5281/zenodo.23063496**

## Author contribution

*G.F.C., B.v.L., B.K., C.V., S.V.A., D.B., E.L., B.S., R.J.V., D.R., E.H.J.K., J.G.C.v.H., H.P.S., and R.C.v.W. wrote the manuscript; G.F.C., B.S., E.H.J.K., J.G.C.v.H., H.P.S., and R.C.v.W., designed the research; G.F.C., B.v.L., B.K., C.V., S.V.A., D.B., E.L., B.S., R.J.V., D.R., E.H.J.K., and R.C.v.W. performed the research; G.F.C., B.v.L., B.K., C.V., S.V.A., D.B., E.L., B.S., R.J.V., D.R., E.H.J.K., and R.C.v.W. analyzed the data*.

## Supplementary material to

