## Supplementary material for "Accelerating drug development using zebrafish disease models supported by pharmacokinetic-pharmacodynamic modeling as new approach methodology (NAM)"

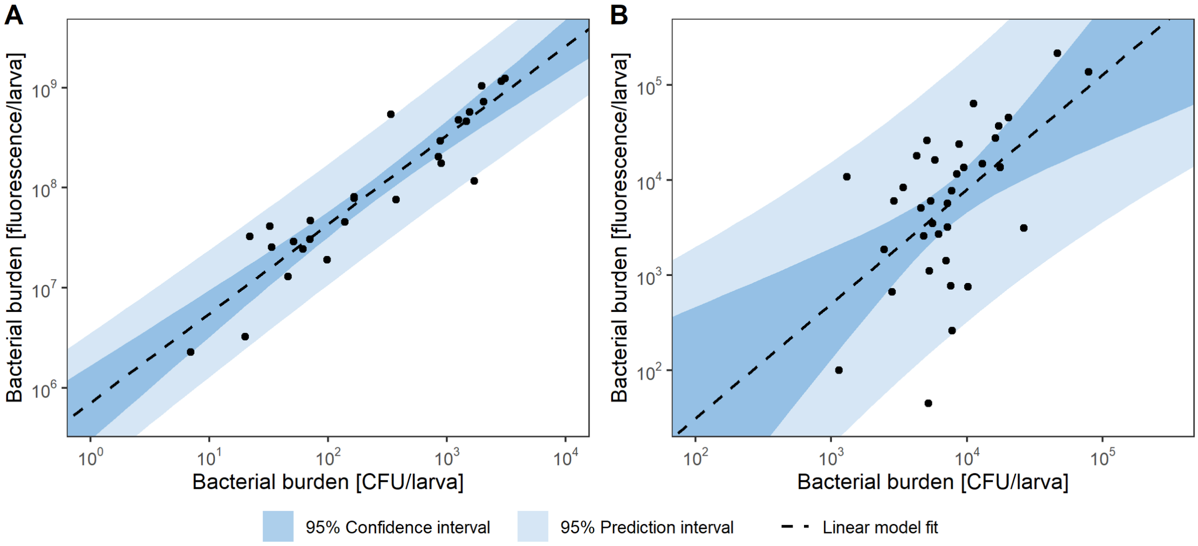


**Figure S1.** Correlation regression including 95% confidence (dark blue) and prediction (light blue) interval between bacterial burden quantified in colony forming units and fluorescent pixel integrated intensity for (A) *Mycobacterium marinum* and (B) *Mycobacterium avium complex*), with an adjusted coefficient of determination ($R^2$) of 0.85 and 0.48, respectively.

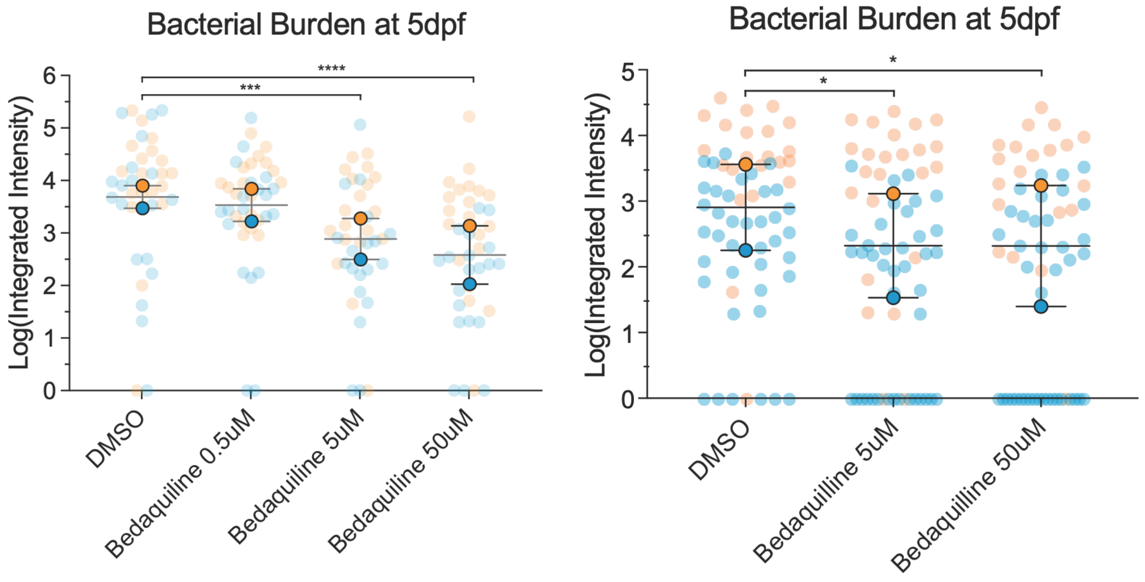


**Figure S2.** Reduction of the bacterial burden of *Mycobacterium avium complex* after 2 days of bedaquiline treatment on (left) yolk-infected embryos and (right) blood-island infected zebrafish embryos/larvae. Colors indicate independent experimental replicates. Each transparent dot is the measurement from one individual; the highlighted points indicate the experimental average. Groups were compared by Kruskal–Wallis with Dunn's post-hoc comparisons against the DMSO control: * $p < 0.05$; ** $p < 0.01$, *** $p < 0.005$; **** $p < 0.001$.

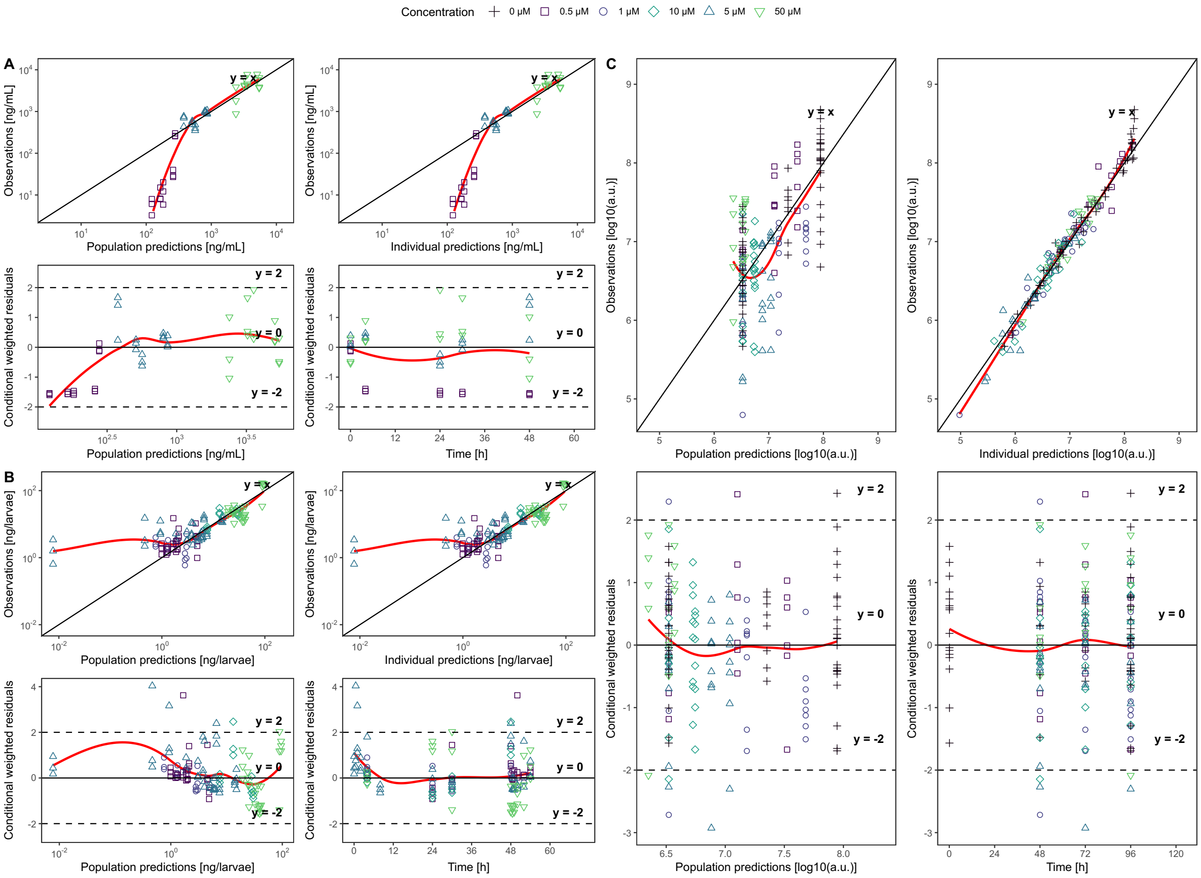


**Figure S3.** Goodness-of-fit plots of the pharmacokinetic and pharmacodynamic models. Shown are the observations, predictions, and residuals for the model fits on the bedaquiline in the medium (A), zebrafish larvae (B), and the bacterial burden (C) in arbitrary units (a.u.).

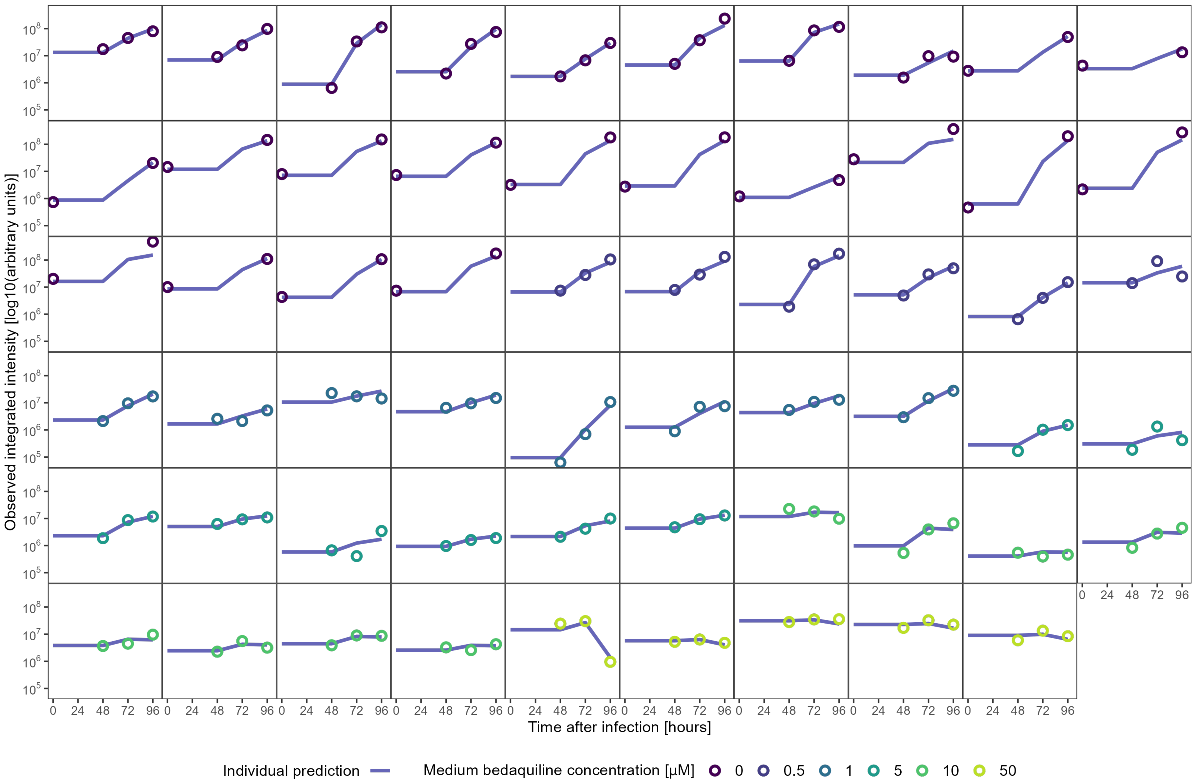


**Figure S4**. Observed and predicted bacterial burden in integrated intensity over time of zebrafish embryos/larvae after infection with *Mycobacterium marinum* and treatment with bedaquiline. Individual panels display data for individual zebrafish embryos/larvae. Open circles represent the observed data points, colored by medium concentration ranging from 0 µM to 50 µM. Solid blue lines show the individual model predictions for each respective zebrafish.

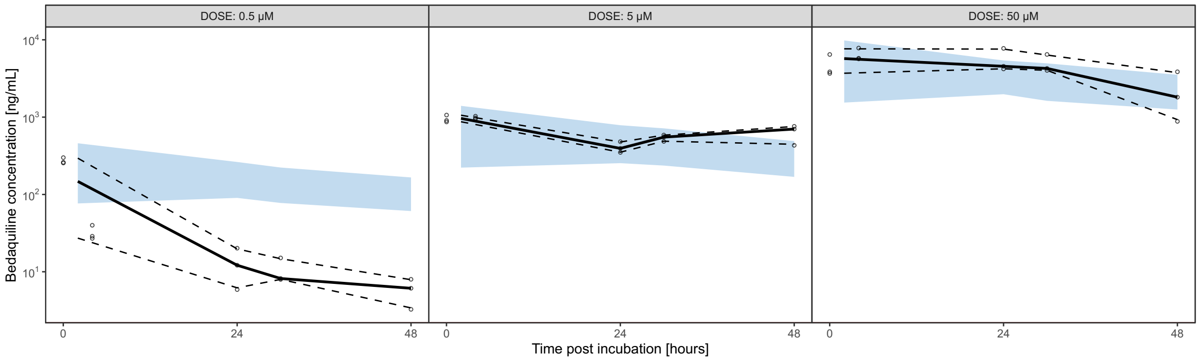


**Figure S5.** Visual predictive checks of simulations (*n* = 500) of the model fit for the medium bedaquiline dynamics. Shown are the individual observations (open circles), the median (solid black line), and the 95% quantile (dashed black) along the 95% prediction interval (blue ribbon). The decrease in the concentration of bedaquiline in the medium when dosed with 0.5 μM was greater than described with the estimated degradation rate constant. Modeling this difference through separate degradation rate constants improved the model significantly but led to unacceptably high relative standard errors in the estimates while resulting in minor changes in pharmacokinetic estimates.

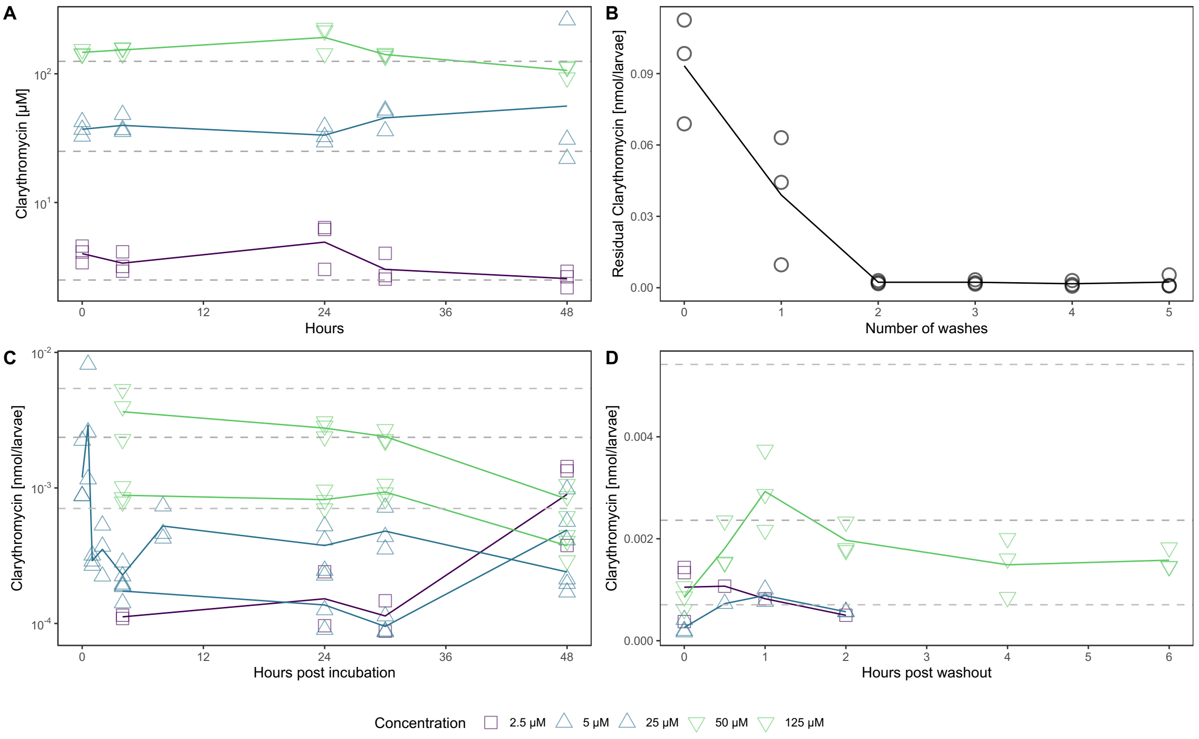


**Figure S6.** Clarithromycin has negligible absorbance across a 2.5-125 µM dose range via waterborne treatment in zebrafish larvae between 3 and 5 days post fertilization. (A) Clarithromycin concentration in the medium during treatment, dotted lines show nominal concentration; (B) residual clarithromycin in washing medium after number of times zebrafish were washed to prevent compound from sticking to the skin; (C) clarithromycin exposure in zebrafish during the constant exposure experiment; (D) clarithromycin exposure in zebrafish during the washout experiment.

### Supplementary methods

**Determination of Maximum Tolerated Dose**

Prior to assessment of drug efficacy in the infection model, the MTD of each test compound on zebrafish larvae between 3 days post fertilization (dpf) and 5 dpf was calculated. MTD was defined as the highest concentration of the compound that caused no significant adverse effects, including mortality, developmental defects, or changes in behavior relative to vehicle controls. All tested compounds were initially dissolved in DMSO to prepare a high-concentration stock solution, and further dilutions in E3 medium were generated to ensure the final DMSO concentration did not exceed 1% (v/v) across all treatment groups.

**Bacterial infections**

Infection experiments were performed according to standard procedures[2,3]. In short, fresh inoculum of *Mycobacterium marinum* (Mm) or *Mycobacterium avium complex 101* (MAC101) expressing the fluorescence protein mWasabi was prepared for every experiment as described previously and resuspended in phosphate-buffered saline (PBS) containing 2% (wt/vol) polyvinylpyrrolidone (PVP40) at ~200 colony forming units (CFU)/nL as determined by optical density (an $OD_{600}$ of 1 corresponds to ~100 CFU/nL). Microinjections were performed using borosilicate glass microcapillary injection needles (Harvard Apparatus, 300038, 1-mm outside diameter [o.d.] × 0.78-mm inside diameter [i.d.]) prepared using a micropipette puller device (Sutter Instruments Flaming/Brown P-97). Needles were mounted on a micromanipulator (Sutter Instruments, MM-33R) positioned under a stereomicroscope and zebrafish embryos between 30 and 32 hours post fertilization (hpf) were injected into the blood island with 1 nL of inoculum using a Picopump (World Precision Instruments, PV820).

**Quantification of drug solubility and stability in E3-buffered medium**

Bedaquiline and clarithromycin were dissolved in E3-buffered medium at 0.25, 1, and 10X MIC concentrations in E3-buffered medium. At 0, 24, and 48 h after incubation at 28.5°C, 1 mL of the medium was collected in a 2 mL Safe-Lock tube (Eppendorf Nederland B.V., Nijmegen, The Netherlands) with the internal standard. All liquid was evaporated overnight using a SpeedVac vacuum. The precipitate was resuspended in 500 mL of 50/50 methanol/water solution and transferred to a Micronic 1.4 mL U-shaped push-cap tube to be analyzed by mass spectrometry.

**Quantification of internal exposure**

Internal exposure of bedaquiline and clarithromycin in zebrafish larvae was quantified as drug amounts in larval homogenates similarly as previously described[2,4–6]. Shortly, groups of n=5 larvae at 3 dpf were treated in 12-well plates containing 2 mL of E3-buffered medium with the desired drug concentration using Netwell inserts (Corning Life Sciences B.V., Amsterdam, The Netherlands). At the time of quantification, larvae were washed and transferred to Safe-Lock tubes. Excess volume was removed, and 100 µL of 90/10 methanol/water solution (v/v) containing internal standard was added after which the larvae were snap frozen in liquid nitrogen and stored at −80°C if needed.

For processing, samples were sonicated at 20% power, 30 seconds pulse, 10 seconds rest on ice. Homogenates were centrifuged at 4000 x g, 4°C for 10 minutes and the supernatant (100 µL) was transferred to a Micronic 1.4 mL U-shaped push-cap tube. Finally, 400 µL of 40/60 methanol/water was added to the sample, thus yielding a final volume of 500 µL of 50/50 methanol/water to be quantified by mass spectrometry.

**Bioanalytical methods**

Quantitation of bedaquiline in zebrafish homogenate samples of the bedaquiline dosed groups was performed by ultra-high performance liquid chromatography linked to mass spectrometry (UPLC-MS) analysis. All samples were injected as received from the sample pretreatment (methanol/water 1/1). Calibration standards for bedaquiline and quality control samples to cover the calibration range were prepared in DMSO. Calibration standards, quality control samples and study samples were processed at the same time. UPLC-MS analysis was carried out on a Triple Quad 6500+ mass spectrometer (AB Sciex, Toronto, Canada), which was coupled to an LC30 AD UPLC-system (Shimadzu, Scientific Instruments, MD, USA). Sample extracts (0.4 µl) were injected onto an UPLC column, (Acquity UPLC BEH C18 1.7µm 2.1*50mm (Waters Corp, Milford, USA)) where bedaquiline and its stable isotope labelled internal standard were separated (retention time 2.66 minutes) from other matrix components using solvents A (0.1 % formic acid in water) and B (methanol) by a linear aqueous gradient from 0-3 minutes 80% -> 10% A, followed by 1 minute cleaning at 2% solvent A and an equilibration step at 80% solvent A for a final minute with a flow-rate of 0.6 ml/min, and a column temperature of 50 °C. The MS was operated in the positive ion mode using electrospray ionization and the parameters were optimized for the quantification of bedaquiline (MRM transition m/z 555.1 > 58) and the stable isotope labeled internal standard (MRM transition m/z 561.2 > 64) using a dwell time of 75 ms per channel and a collision energy of 71 eV. A log-log transposed linear regression model was used to calibrate peak area ratios of the analyte to its internal standard against the analyte concentrations. The concentrations of the analyte in the study samples were calculated by interpolation from the calibration curve using Analyst Software (version 1.7.1) from AB Sciex, and normalized by the number of larvae in each sample to result in the total internal amount per larva (ng/larva), which was divided by the total volume of the larva to reach internal concentration (ng/mL)[4,6] .

#### Model code

```
; Description:  PKPD Model for zebrafish larvae infected with Mycobacterium marinum
and treated with bebaquiline via waterborne exposure.
; Author:          G. Forn-Cuni, B. van Lieshout
, Elke Krekels

;--------------------------------------------------------------------
; Columns and Units
;--------------------------------------------------------------------
; ID
; TIME       Hours post treatment
; AMT        Concentration dosed (ng)
; CMT        1 EW  2 Larvae
; DV         Amount measured.
;                For CMT1: ng/mL
;                For CMT2: total ngs in larvae
;                For CMT3: Integrated fluorescence intensity
; Grouping variables:
; DOSE        The grouping variable for drug concentrations
; DPF        Days post fertilization

;--------------------------------------------------------------------
; Modeling
;--------------------------------------------------------------------
$PROBLEM PKPD
$INPUT ID AMT DOSE DV MDV EVID CMT TIME DPF
$DATA data.csv IGNORE=@

$SUBROUTINE ADVAN13 TOL=9

$MODEL
COMP                ; CMT 1 PK - Bedaquiline amount in EW
COMP                ; CMT 2 PK - Bedaquiline amount in Larvae
COMP                        ; CMT 3 PD - Bacterial Burden

$PK
;Typical values, scaled
TVK10               = THETA(1)/1e2          ; Elimination rate from CMT1.
Evaporation, protonation, etc.
TVDCF1              = THETA(2)              ; Correction factor 0.5< <=10
TVDCF2              = THETA(3)              ; Correction factor 10<
TVK12               = THETA(4)/1e3          ; Rate constant of absorption in larvae
TVK21               = THETA(5)/1e1          ; Rate constant of elimination in larvae
TVK21_5dpf          = THETA(6)              ; Increase in clearance at 5 dpf

TVKGROWTH           = THETA(7)/1e1      ; Rate of bacterial replication
```

```
TVINOC              = 10**THETA(8)       ; Inoculum estimate
TVBMAX              = 10**THETA(9)       ; Carrying capacity
TVTLAG_KG           = THETA(10)          ; Delay before growth

TVEMAX              = THETA(11)          ; (Maximum) effect size
TVEC50              = THETA(12)*1000     ; Concentration at 50% effect size
TVEMAX_3dpi         = THETA(13)          ; Increase in effect at 3 dpi

;Individual values
K10                 = TVK10      * EXP(ETA(1))
DCF1                = TVDCF1     * EXP(ETA(2))
DCF2                = TVDCF2     * EXP(ETA(3))
K12                 = TVK12      * EXP(ETA(4))
K21                 = TVK21      * EXP(ETA(5))
K21_5dpf            = TVK21_5dpf  * EXP(ETA(6))

KGROWTH             = TVKGROWTH   * EXP(ETA(7))
INOC                = TVINOC     * EXP(ETA(8))
BMAX                = TVBMAX     * EXP(ETA(9))
TLAG_KG             = TVTLAG_KG   * EXP(ETA(10))

EMAX                = TVEMAX     * EXP(ETA(11))
EC50                = TVEC50     * EXP(ETA(12))
EMAX_3dpi           = TVEMAX_3dpi * EXP(ETA(13))

IF(DPF.GT.4)K21 = K21 + K21_5dpf
IF(DPF.GE.4)EMAX = EMAX_3dpi

;Compartments
F1 = 1
IF (DOSE.GT.0.5) F1 = DCF1            ; DCF in CMT1. Sticking to plastic, etc.
IF (DOSE.GT.10) F1 = DCF2             ; Precipitation at highest concentrations

V1 = 2                  ; Volume from CMT1, fixed at 2mL (DVs in ng/mL, AMT in ng)
S1 = V1                               ; Scale concentrations to total amounts in CMT1
S2 = 1                                ; ZF units are ng already, so no need to scale CMT2

A_0(3) = INOC                 ; Initial inoculum estimate set as compartment initial value

VOLUME = 0.000253             ; Larval volume
IF (DPF.EQ.4) THEN
  VOLUME = 0.000263           ; Larval volume
ENDIF
IF (DPF.EQ.5) THEN
  VOLUME = 0.0003             ; Larval volume
ENDIF
```

```
$DES
; PK
DADT(1) = - K10 * A(1) - K12 * A(1) + K21 * A(2)
DADT(2) = K12 * A(1) - K21 * A(2)

; PD
C2 = A(2)/VOLUME          ; Convert internal amounts to concentration based on larval volume for PKPD
EFF = 0             ; The effect when there is no concentration or NA is 0.

IF(A(2).GT.0) THEN
  EFF = (EMAX * C2) / (EC50+C2)    ; Emax effect
ENDIF

KG = KGROWTH * (1-(A(3)/BMAX))   ; Logistic

IF(T.LT.TLAG_KG) THEN
KG = KG * 1e-6 ; Small number to keep IIV working
ENDIF

DADT(3) = KG * A(3) * (1-EFF) ; Proportional drug effect

$ERROR
IPRED = F
IF (CMT.EQ.1) THEN
Y = F * (1 + EPS(1))
ENDIF

IF (CMT.EQ.2) THEN
IPRED = A(2)
Y = IPRED * (1 + EPS(2)) + EPS(3)
ENDIF

IF (CMT.EQ.3) THEN
IPRED = LOG10(A(3)+0.00001)
Y = IPRED + EPS(4)
ENDIF

; Variables for reporting
A1 = A(1)
A2 = A(2)
A3 = A(3)
C_eff = A2/VOLUME

$THETA
(1.89)  FIX ;K10 [1/h] 1e-2
(0.299) FIX ;F1 >0.5 [-]
```

```
(0.216) FIX ;F1 >10 [-] 1
(0.941) FIX ;K12 [1/h] 1e-3
(0.89)  FIX ;K21 [1/h] 1e-1
(0.122) FIX ;K21_5df [-]
(0.853) FIX ;KGROWTH [1/h] 1e-1
(6.52)  FIX ;INOC [RFU] 10^x
(8.19)  FIX ;BMAX [RFU] 10^x
(48)    FIX ;T_LAG KG [h] 1
(1.09)  FIX ;EMAX [-] 1
(26.5)  FIX ;EC50 [ug/mL]
(1.37)  FIX ;EMAX_3dpi [-] 1

$OMEGA 0     FIX  ; K10
$OMEGA 0     FIX  ; F1 >0.5
$OMEGA 0     FIX  ; F1 >10
$OMEGA 0     FIX  ; K12
$OMEGA 0     FIX  ; K21
$OMEGA 0     FIX  ; K21_5dpf
$OMEGA 0.442 FIX  ; KGROWTH
$OMEGA 1.5   FIX  ; INOC
$OMEGA 0     FIX  ; BMAX
$OMEGA 0     FIX  ; T_LAG KG
$OMEGA 0     FIX  ; EMAX
$OMEGA 0     FIX  ; EC50
$OMEGA 0     FIX  ; EMAX_3dpi

; Residual variability
$SIGMA 0.381  FIX  ; EPS1 CMT1 prop
$SIGMA 0.247  FIX  ; EPS2 CMT2 prop
$SIGMA 11.1   FIX  ; EPS3 CMT2 add
$SIGMA 0.0443 FIX ; EPS4 CMT3 add (log10)
```
